# Urban colonization is driven by a mixture of evolutionarily conserved and labile traits

**DOI:** 10.1101/2020.01.20.912170

**Authors:** David A. Duchene, Carolina Pardo-Diaz, Maider Iglesias-Carrasco

## Abstract

Urbanization is a fast and dramatic transformation of habitat that generally forces native fauna into novel ecological challenges. The biological prerequisites necessary to establish in urban areas have been widely studied, but the macroevolutionary characteristics of traits that allow urban colonization remain poorly understood. Urban colonization might be facilitated by traits that are evolutionarily conserved and which lead to a diversity of closely related species. Alternatively, urban colonization might be associated with labile traits that frequently arise and are lost. In a large data set from passerine birds, we find that urban colonization has a signal of highly labile traits, despite many traits associated with colonization being highly conserved. Urban colonization is associated with traits that allow faster speciation than non-urban-colonizing counterparts, and more frequently transition to non-urban trait states than in the opposite direction. Overall, the traits that facilitate urban colonization are a mix of highly conserved and labile traits and appear to provide an evolutionarily successful strategy.

## Introduction

Urbanization is one of the most rapid anthropogenic changes to the environment, and can have a dramatic impact on local communities [1,2]. While urbanization is often associated with a loss of biodiversity and of evolutionary distinctiveness [3], a broad range of species have successfully established in these novel environments. Understanding the biological traits that allow urban colonization can be important for predicting the impact of urbanization on biodiversity. In birds, there are several biological predictors of successful urban colonization including lack of plumage dichromatism, large body mass, large brain size, migration ability and broad environmental tolerance [4–8]. Such a collection of traits might impact species survival, leaving a mark across deep evolutionary timescales [9,10]. However, the macroevolutionary trajectory of the complete set of traits in urban-colonizing species remains poorly understood.

The macroevolution of traits that facilitate urban colonization is driven primarily by three processes: speciation, extinction, and the transition among species that do not have such traits and those that do. In consequence, the distribution of urban colonizing species in a phylogeny will depend on the rate of each of these processes. One possible scenario is that the transition between traits that do not allow urban colonization and those that do is infrequent in either direction. In such cases, the traits that allow colonization might only occur among closely related species and will be clustered across the phylogeny. Any differences in the current diversity of the urban colonizers versus non-colonizers is then driven by differences in rates of speciation and extinction, and not by the rate of appearance and disappearance of traits. In birds, however, urban colonizers come from a broad range of taxa, suggesting that the rate of transition between traits is actually high [3,11].

In another scenario, species might frequently transition between having traits that facilitate urban colonization and those that do not. The speciation and extinction rates of urban colonizers might be similar to those of non-colonizers, but the traits that allow colonization might be frequently gained and lost. If urban colonization is driven by such evolutionarily labile traits, colonizers and non-colonizers will be highly dispersed across their phylogeny. However, there is also evidence against this hypothesis, since many of the traits associated with urban colonization are conserved, such as brain size [11].

Alternatively, urban-colonizers and non-colonizers might differ in rates of both diversification (speciation and extinction) and transition. In this case, colonizers and non-colonizers might differ in their pattern of phylogenetic clustering, and several combinations of rates of speciation, extinction and transition can lead to such an observation. One interesting case occurs when transition is high in a single direction (e.g., non-urban colonizer traits to urban-colonizer traits), and the frequently-emerging traits cause an increased rate of extinction by benefitting individuals at the expense of the population. This phenomenon has been termed “evolutionary suicide” [12,13]. Examples of traits that have been proposed to emerge often but reduce the evolutionary success of species include self-compatibility in plants [14] or some forms of specialization [15]. The traits that allow urban colonization might be such an evolutionary “dead-end”, if urban colonization increases the already-high chances of extinction in those species. This hypothesis might explain the loss of evolutionary distinctiveness in urban areas [2] and the lack of phylogenetic clustering in urban colonizers [3], and should appear as a highly dispersed or “tippy” pattern of urban colonizers in their phylogeny [16].

Apart from the main three scenarios explained so far, there are other combinations of rates of speciation, extinction, and transition that might govern the macroevolution of current urban-colonizers and non-colonizers. Therefore, to test the pattern associated with urban colonization and whether it is driven by traits that are evolutionarily successful or detrimental, and either conserved or labile, we analyzed published data from passerine birds. We first tested whether urban colonization and three traits previously associated with urban colonization (brain size, body mass, and plumage dichromatism) are phylogenetically dispersed within the passerines. We then used maximum likelihood and simulation approaches to test whether the collection of traits that might have allowed colonization of urban areas are either evolutionarily conserved, or tend to be either labile or even suicidal at a macroevolutionary scale.

## Methods

To gain insight into the types of traits that allow urban colonization, we examined the macroevolutionary features of the species that have successfully colonized urban areas. We gathered information about urban presence/absence [17] of passerine birds, and three traits of species associated with the colonization and establishment of passerines in urban areas: brain size ([4,18], but see [11,19]), body mass [5,8] and plumage dichromatism [8] (see Supplementary Information for data collection).

We first tested whether brain size, body mass, plumage dichromatism and urban colonization of the set of species for which the latter variable was available are either taxonomically grouped or randomly dispersed across their phylogeny. Previous studies have already explored the phylogenetic characteristics of brain size, body mass, plumage dichromatism in datasets including large samples of the extant passerine species (e.g., [20]). To explore whether similar patterns are seen only for the set of species for which there are data on presence/absence in urban areas, we ran these analyses on the reduced data set for which all variables were available (brain size, n = 251; body mass, n = 506; plumage dichromatism, n = 506).

We first estimated the maximum clade credibility (MCC) phylogenetic tree of a random sample of 1000 time-calibrated trees containing every species of birds [21]. The complete MCC tree was pruned to contain the 506 species of passerine birds for which the other data were available (or 251 species for the dataset in brain size), and was then used for subsequent analyses. We tested whether urban habitat colonization is either significantly phylogenetically clustered or over-dispersed by calculating Fritz and Purvis’ *D* statistic [22] and comparing that value to 1000 simulations under each of the two null models, using the R package *caper* [23]. Similarly, we tested the hypotheses that the variables of brain size, body mass and plumage dichromatism have phylogenetic signal by testing the significance of Pagel’s *λ* [24] as implemented in the R package *phytools* [25].

To assess whether traits that allow urban colonization are highly labile or even suicidal, we performed two tests of the macroevolution of successfully colonizing species (see [16] for full details on this method): (i) we tested whether colonizing species have had a greater number of evolutionary origins (*NoTO*), and (ii) whether they have a lower number of also-colonizing sister species (*SSCD*) than expected under a null process in which rates of gain and loss of a trait are equal in both directions (as proposed in [16]), using the R package *phylometrics* [26]. Traits that are likely to be significantly labile or suicidal have values of these metrics that are significantly different from the null, and also have a *D* statistic of phylogenetic clustering indicating a significantly dispersed phylogenetic distribution [16]. We performed these analyses assuming that we have an incomplete sample from the passerines (506 / 5966 = 0.085).

We examined whether the data contained a signal of significant differences in rates of speciation, extinction, or transition between colonizing and non-colonizing species by testing models of binary state-speciation and extinction (BiSSE; [27,28]). The diversification rates in passerine birds are likely to be affected by many factors that are not considered in this study (e.g., [29]). Therefore, we restricted our analyses to testing whether the distinction between colonizers and non-colonizers is superior to more simple models where the distinction is not present. If the primary drivers of diversification rates in passerines are entirely unrelated with the traits associated with urban colonization, the distinction between colonizers and non-colonizers will not be meaningful and lead to similar fit in models of diversification.

We tested whether a model in which colonizing and non-colonizing species have different rates of speciation, extinction, and transition (hereafter “full model”), was superior to four alternative models in which colonizing and non-colonizing species have (i) identical speciation rates, (ii) identical extinction rates, (iii) identical transition rates, or (iv) equivalent values for each of these traits across the two groups. The models were corrected for incomplete taxon sampling and likelihood ratio tests were performed comparing the full model and each of these alternative hypotheses. Results were corrected for multiple tests using false discovery rates.

## Results

We found that in passerine birds there is significant phylogenetic signal in plumage dichromatism *(λ* = 0.78, *p* < 0.001), brain size *(λ* = 0.88, *p* < 0.001), and body mass *(λ* = 0.88, *p* < 0.001). Meanwhile, colonization of urban areas is significantly more phylogenetically dispersed than a pattern emerging from Brownian motion. However, it is also significantly more clustered than a random assortment of species across the phylogeny of passerine birds (*D* = 0.695, *p* < 0.001; Fig. 1). Nonetheless, we find that evolutionary factors associated with urban colonization lead to a greater number of evolutionary origins, and a smaller number of also-colonizing sister taxa than expected under a Brownian motion evolutionary process (*NoTO* = 3.35, *p* < 0.001; *SSCD* = 193.21, *p* < 0.001).

**Figure 1.**
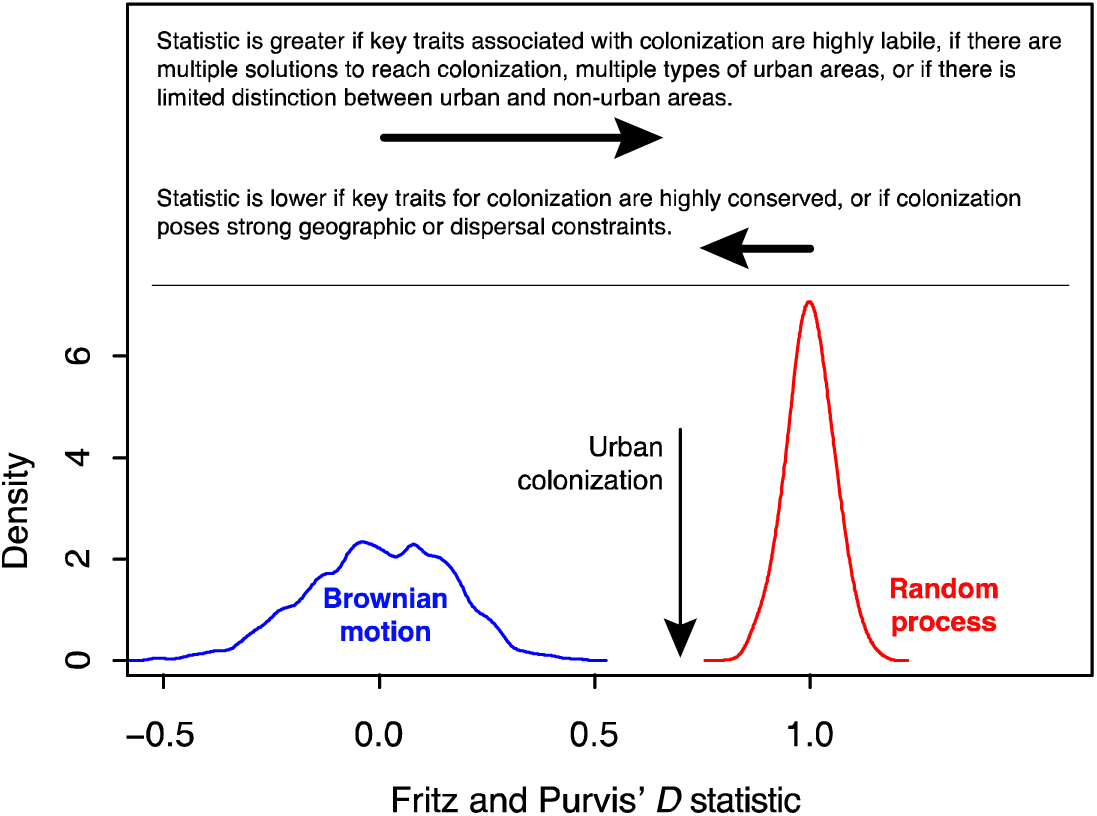
Fritz and Purvis’ *D* statistic of phylogenetic clustering for urban colonization (black vertical arrow) as a proxy of the traits that allow this phenomenon. Expected distributions shown are drawn from simulations of the data under an evolutionary processes of Brownian motion (blue) and random change (red). We describe some of the factors that might cause an increase in phylogenetic dispersal versus clustering of traits associated with urban colonization.

**Figure 2.**
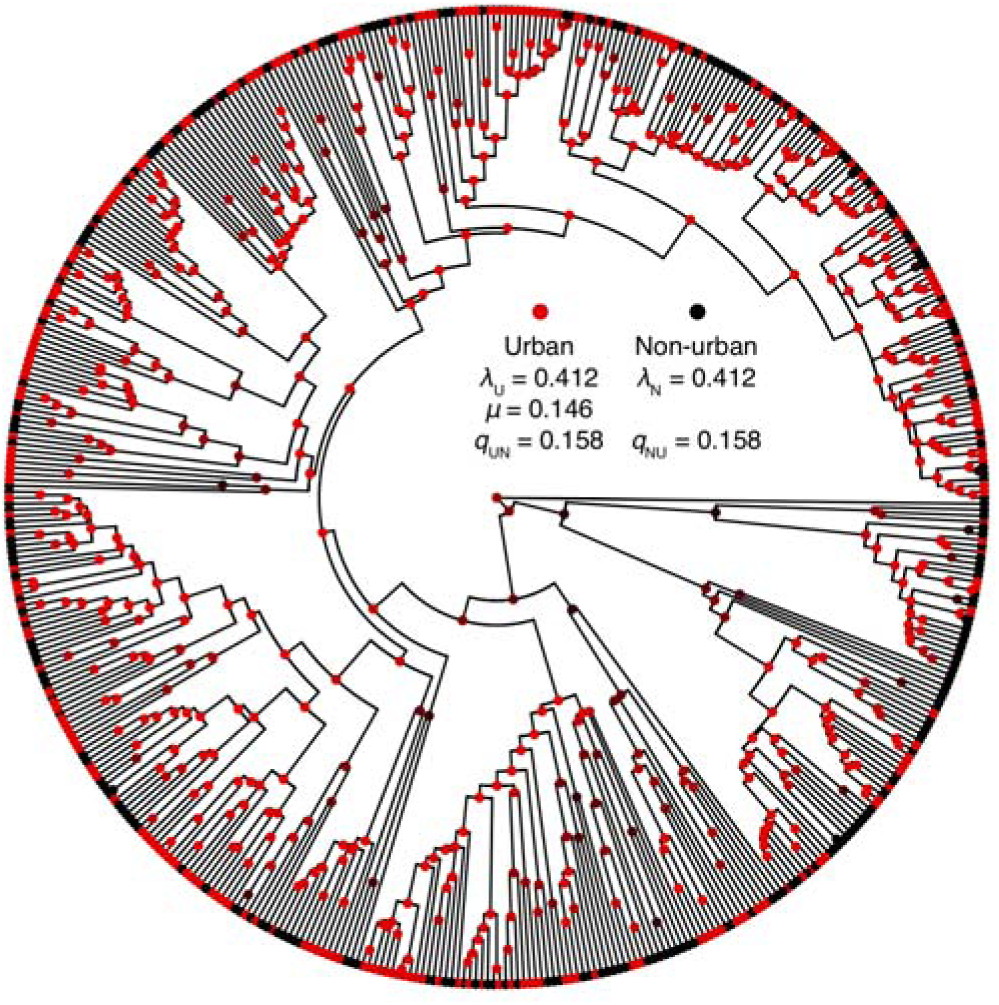
Phylogenetic tree of the passerine species sampled showing the positioning of urban colonizers (red) and non-urban colonizers (black). Internal nodels and the statsitics shown are drawn from the maximum likelihood estimate of the ancestral states under the BiSSE model. The estimates shown are not intended to represent the model that best describes the processes the drove the diversification of passerine birds.

Tests of BiSSE models showed that the full model, which makes a distinction between urban colonising species and non-colonizers, provided a significantly better fit than models that did not make a distinction between the two groups in terms of their speciation rates (*p* = <0.001), rates of transition (*p* = 0.003), or in all of speciation, extinction, and transition rates (*p* < 0.001; Fig. 1; Table S1). However, the full model was not significantly superior to a model in which the two groups share an identical extinction rate (*p* = 0.099; *ε* = 0.146). In the full model and the more simple model where extinction rates are identical, speciation rates were estimated to be greater in urban colonizing species *(λ*_U_ = 0.412) than in non-colonizing species *(λ*_N_ = 0.149), and the estimated rate of transitions between states was estimated to be greater in the direction from colonizing species to non-colonizers (*q*_UN_ = 0.158) than the opposite (*q*_NU_ = 0.023).

## Discussion

Some collections of traits can influence the chances of species success or demise, so the macroevolutionary history of different groups of traits have been a longstanding matter of interest (e.g., [10,22]). Our data show that rates of transition and speciation might be uneven while extinction rates are similar. This suggests that many of the traits that are associated with urban colonization of species of passerine birds are evolutionarily conserved. However, urban colonizers are not significantly grouped phylogenetically in passerine birds, suggesting that urban colonization is not driven by purely conserved or labile traits. Some of the traits that allow urban colonization are likely to be labile and have frequently been lost in the evolution of passerines. Interestingly, the complete collection of traits has provided a comparatively successful strategy over the traits that occur in non-urban species. Therefore, our data suggest that urban areas are not a sink of biodiversity that accelerates the demise of species with already unfortunate macroevolutionary trajectories.

Some of the traits explored here might have direct effects on the elevated rate of speciation in urban-colonizing species. However, this effect occurs in the opposite direction as might be expected, because urban species tend have large body size and have lower amounts of sexual selection [8], both of which are associated with slower rates of diversification [20,30]. Instead, other features of urban colonizers might have been associated with their relatively high speciation rates. One example is being a dietary generalist, which is strongly associated with diversification rates in birds [20] and is a distinctive feature of urban colonizers [11].

Urban colonizers are associated with relatively fast speciation rates and frequent transition towards a different combination of traits. Some successful evolutionary strategies can frequently breakdown and lead to novel diversity. Examples include the common switch from self-incompatibility to the less-successful self-compatibility strategy in the plant family Solanaceae [14], or the frequent switch from bisexuality to the less-successful unisexuality in liverworts [31]. Similarly, generalist taxa sometimes have faster rates of diversification and greater rates of transition to specialization than the opposite direction [15]. In addition to being dietary generalists, urban-colonizing species have been proposed to have broad environmental tolerances [7] and high behavioural flexibility ([4,32] but see [11,19]), which are likely easily lost evolutionarily, yet might provide species with resilience and long term evolutionary success.

Our results suggest that urban areas attract species with traits that are robust in macroevolutionary terms, such that cities might serve as a reservoir of biodiversity [33,34]. This is consistent with the lack of evidence for poor health in urban taxa [35–37] and the fact that urban species are often in the process of becoming reproductively isolated from non-urban counterparts [33]. The high rate of transition out of the traits that allow urban colonization means that novel diversity arising from urban areas might contain non-urban traits (e.g., strong sexual selection, small body size, specialization). Understanding the medium- and long-term evolutionary trajectories of urban species might aid urban planning in the future, since the predicted increase in urbanization might impact species without suitable traits (e.g., specialist species).

While urban areas are associated with the homogenization of biodiversity and the loss of evolutionary distinctiveness, the traits that allow for colonization of these areas do not appear to cause increased extinction rates at a macroevolutionary scale. Since urban colonization is not phylogenetically clustered, it is also likely to be driven by highly labile and common traits. Labile traits that are most likely to contribute to urban colonization might include nesting location [5] or song type [38], while other conserved traits might include feeding habits or brain size [4]. Exploring the macroevolution of traits further on species colonization of novel habitats is likely to bring important insights into species sensitivity to future urbanization.

## Author contributions

DAD, MIC, and CPD conceived the idea, MIC and CPD collected the data, DAD analysed the data, all the authors interpreted the results and helped drafting the manuscript.

## Funding

This work was supported by funding from the Australian Research Council to D.A.D. (grant DE190100544).

## Conflict of interest

The authors declare no conflict of interest

## Ethics

Not applicable

## Data accessibility

The data used in this project are available at github.com/duchene/urbanSexSel.

## Notes

https://github.com/duchene/urbanSexSel

