## Supplementary Information for "Urban colonization is driven by a mixture of evolutionarily conserved and labile traits"

**Supplementary methods**

Urban tolerance of species was collected based on published data on bird communities of both urban areas (characterized by >50% of the surface built and >10 buildings per ha) and the nearby natural areas [1]. In the original paper [1], studies were selected by performing an exhaustive literature search using the combination of the terms ‘bird*’, ‘assemblages’ and ‘urban*’ in Web of Science, Google Scholar, and SmartCat. The studies selected had information on the avian communities of urban and non-urban habitats from the same area. Studies were excluded if they did not include the full list of species sampled in both urban and nonurban environments.

This searching criteria ensures covering all the previously published studies in the topic and a good representation of species of urban-colonizing passerines. The list also represents the species that have the potential to invade cities (e.g., they inhabit the surrounding areas), and discards species that live in remote areas far from cities, such that due to dispersal constraints are unlikely to contact urban environments. Therefore, a species was considered to be “absent” from urban areas when it was not found in urban areas but appeared in the nearby non-urban environment. We considered a species to be “present” if it was found in urban areas, independently of its presence or absence in the nearby non-urban environment. The search criteria led to 26 studies from 17 countries in 4 continents and identified 665 bird species from paired urban versus nonurban environments [2]. The information on bird assemblages was standardized by using only community data collected by maintaining a constant field method and time period. We used passerine birds due to their higher representation in urban environments and the abundance of data from the literature, and discarded the non-passerine species from the original dataset. Our data collection led to a data set with information on urban presence/absence for 506 passerine birds.

In order to explore the phylogenetic distribution of specific traits associated with the colonization and establishment in urban areas, we also collected data on three biological traits related to the presence of birds in cities: brain size, body mass and plumage dichromatism. Brain size has been widely suggested to be a strong predictor of the ability of birds to colonize novel habitats, including urban areas ([3,4], but see [5,6]). Brain size was collected from [7] and [6], and measured as brain mass from dead and museum specimens. For statistical analyses we used the relative brain size calculated as the residuals of the regression of the log transformed brain size on log transformed body mass. The dataset of brain size used to calculate the phylogenetic signal of this trait included the 251 species for which we had information in both brain and presence/absence in urban areas.

Body mass has a positive effect on the ability of urban colonization in passerines [8,9]. In general, large-bodied species are more likely to appear in urban environments, probably due to the higher dispersal and competitive abilities, and environmental tolerance [10,11]. Body mass was obtained from the same dataset as urban tolerance from [1], that originally collected the data from the “Handbook of the birds of the world alive” [12]. The dataset included 506 passerine bird species, the same for which we had presence/absence in urban areas (Present in urban areas N = 258 sp; absent from urban areas N = 248 sp).

We used the same dataset of plumage dichromatism used in a recent study [9]. In this previous study, we used plumage dichromatism as a proxy for the strength of sexual selection, and showed that species with lower dichromatism are more likely to be present in urban areas [9]. Plumage dichromatism was calculated as the difference in plumage colour between males and females (score male – score female), using the scores reported in ([13]; see also [9] for more details on the appropriateness of this measurement). Coloration is measured in the original publication for all passerine (N = 5983 species), using spectrometry and digital scans of the images in the “Handbook of the Birds of the World” [14]. A colour plumage score was calculated for males and females from multiple sections that differ between sexes in dimorphic species. Each species had its own measurement, such that the score is independent on the relative coloration measured for other species. The dataset used in this study to calculate the phylogenetic signal of plumage dichromatism included 506 passerine birds, the same for which data of urban presence/absence were available (present in urban areas N = 258 species; absent from urban areas N = 248 species).

**Table S1.** Results from BiSSE models tested, including parameter estimates. Model comparison results show likelihood ratio tests comparing each model to the full model, with *P*-values corrected using false discovery rates.

|  | *λ*_0_ | *λ*_1_ | *μ*_0_ | *μ*_1_ | *q*_01_ | *q*_10_ | D.F. | log likelihood | AIC | χ^2^ | *P*-value |
| --- | --- | --- | --- | --- | --- | --- | --- | --- | --- | --- | --- |
| Full model | 0.116 | 0.506 | 0.074 | 0.274 | 0.041 | 0.134 | 6 | -2126.209 | 4264.418 | - | - |
| Equal speciation rate | 0.391 | 0.391 | 0.076 | 0.461 | 0.245 | 0.040 | 5 | -2152.232 | 4314.463 | 52.045 | <0.001 |
| Equal extinction rate | 0.149 | 0.412 | 0.146 | 0.146 | 0.023 | 0.158 | 5 | -2127.806 | 4265.611 | 3.194 | 0.099 |
| Equal transition rate | 0.115 | 0.768 | 0.000 | 0.614 | 0.108 | 0.108 | 5 | -2131.136 | 4272.271 | 9.854 | 0.003 |
| Base model (all rates equal) | 0.485 | 0.485 | 0.370 | 0.370 | 0.210 | 0.210 | 3 | -2164.420 | 4334.839 | 76.422 | <0.001 |
